# Dihydroartemisinin inhibits Epstein-Barr virus reactivation and replication targeting lytic proteins: insights for drug repurposing

**DOI:** 10.64898/2026.05.25.727607

**Authors:** Hiteshi Vaidya, Manoj Kumar

**Author notes:** Authors’ Email ID.

## Abstract

Epstein-Barr virus (EBV) is an oncogenic virus which is responsible for various malignant as well as non-malignant diseases and leads to about 200,000 deaths each year. Despite efforts, there are no FDA-approved drugs targeting EBV. Reactivation of EBV plays a critical role in the transition from latency to lytic cycle, leading to viral replication and disease progression, and is primarily regulated by the transactivator BZLF1. In this study, we combined computational screening with experimental validation to identify repurposing drugs that inhibit EBV reactivation and replication. FDA-approved compounds predicted using in-house AI/ML-based model (Anti-EBV) and miRNA-seq and RNA-seq analyses, were selected for further evaluation. Molecular docking against BZLF1, supported by *in silico* alanine scanning to identify critical DNA-binding residues, led to the selection of seven candidate drugs. Among these, an antimalarial drug, dihydroartemisinin (DHA), showed the strongest inhibitory activity *in vitro*, with an IC_99_ of 1 µM and an SI Index of 113.5. DHA reduced both EBV viral copy number and the expression of early and late lytic genes. Molecular docking and simulation studies demonstrated stable binding of DHA within the BZLF1 DNA-binding pocket, inhibiting the key residues involved in BZLF1 activation and DNA binding. Analysis at the gene level confirmed its inhibitory effect on EBV replication, while expression analysis at the transcriptional and protein levels, along with immunofluorescence analysis, indicated its inhibitory effect on EBV reactivation and virion assembly. These findings suggest DHA as a promising repurposing antiviral candidate targeting EBV lytic proteins and offers an effective target-based therapeutic strategy.

**Importance:** This study identifies a repurposed small-molecule inhibitor of EBV reactivation and replication. Here, we proposed target-based therapy, integrating computational and experimental approaches to target the EBV lytic transactivator BZLF1. Since early lytic EBV protein BZLF1 plays a critical role in viral reactivation and replication, inhibition of its activation and DNA-binding function represents a promising therapeutic approach to prevent EBV infection. Molecular docking and simulation studies revealed stable binding of DHA within the BZLF1 DNA-binding pocket. Furthermore, in vitro analyses demonstrated significant inhibition of viral gene copy number and reduced mRNA and protein levels of key lytic proteins. Thus, this study demonstrated DHA as a safe and effective repurposed therapeutic candidate against EBV infection.

## Introduction

Epstein-Barr virus (EBV), also known as Human herpesvirus 4 (HHV-4), is an oncogenic virus ^1^. It is a double-stranded DNA virus with a genome size of approximately 172 kb and belongs to the genus Gammaherpesvirus (subgenus Lymphocryptovirus) under the family Herpesviridae^2^. EBV infects more than 95% of the global population^3^ and is globally associated with nearly 240,000 to 360,000 cancer cases and approximately 200,000 deaths each year^4^. EBV is linked to a wide range of malignancies, including gastric carcinoma, nasopharyngeal carcinoma, Burkitt lymphoma, Hodgkin lymphoma, non-Hodgkin lymphoma, and diffuse large B-cell lymphoma^5^. In addition, it is associated with non-malignant conditions such as infectious mononucleosis, multiple sclerosis, and oral hairy leukoplakia ^6,7^. EBV transmission primarily occurs through saliva, including sharing drinks and the consumption of pre-chewed food. Less common routes, such as blood transfusion, stem cell transplantation, and organ transplantation, have also been reported^8,9^. Although EBV mostly remains dormant in infected cells, reactivation into the lytic cycle is required to replicate and spread^10^. This transition plays an important role in disease progression, as it leads to the production of new viral particles. EBV reactivation is initiated by two immediate-early genes, BamHI Z fragment leftward open reading frame 1 (BZLF1) and BamHI R fragment leftward open reading frame 1 (BRLF1). BZLF1 is a DNA binding protein that reactivates latent infection and alters the structure of the viral genome, making it accessible for transcription^11^. BRLF1 increases transcription of downstream lytic genes, triggering a cascade that leads to complete lytic activation^12–14^. Compared to BRLF1, BZLF1 plays a central role in EBV lytic reactivation. Upon phosphorylation at the S186 residue, BZLF1 activates the expression of both the BZLF1 and BRLF1 genes through promoter binding, thereby disrupting viral latency^11,15^. In addition, it promotes viral DNA replication by binding to the oriLyt region via its DNA-binding domain^16^. Since BZLF1 play a significant role in virus reactivation, it is an important therapeutic target, and its inhibition may block viral reactivation and limit disease progression.

Most studies aimed at identifying antiviral agents for EBV have mainly targeted DNA replication. These include nucleoside analogues^17^, plant-derived compounds ^18^, and small organic molecules^19^. Furthermore, there are limited studies targeting EBV reactivation^20,21^. Despite efforts, no drugs have yet been approved by the Food and Drug Administration (FDA) or the European Medicines Agency (EMA) for the treatment of EBV-associated diseases^22^. Hence, there is a need to identify effective, non-toxic antivirals to prevent viral reactivation and replication.

Drug repurposing has become a promising approach in recent years to identify new therapeutics by repurposing approved drugs with known safety profiles^23^. This approach significantly reduces the time, cost, and risk associated with traditional drug development and helps in faster translation into clinical applications.

In the present study, FDA-approved drugs from the DrugBank database were screened using two *in silico* approaches to identify potential repurposing drugs against EBV, which were subsequently tested *in vitro* for antiviral activity. FDA-approved drugs obtained from an in-house AI/ML-based predictive algorithm, Anti-EBV^24^, and *in silico* miRNA-Seq and RNA-Seq analysis of differentially expressed genes in EBV-infected cells were selected for further screening^25^. Since BZLF1 plays an important role in EBV lytic reactivation, targeting its activation and DNA-binding activity may be a promising therapeutic strategy. Therefore, critical DNA-binding residues of BZLF1 were first identified using *in silico* alanine scanning.

Subsequently, the selected drugs were docked to this key EBV lytic protein to evaluate their binding affinity and inhibitory potential. Based on docking results, seven potential drugs binding at the DNA binding domain of BZLF1 were selected for experimental validation in the EBV-positive P3HR-1 cell line. Among all the drugs, Dihydroartemisinin (DHA), a derivative of the Chinese herb with high efficacy and low toxicity known for its antimalarial effect and antitumor activity ^26,27^, demonstrated the most significant inhibitory effect on EBV reactivation and replication. Since early lytic EBV protein BZLF1 is critically involved in viral reactivation and replication, targeting it can be a promising therapeutic approach to prevent EBV infection.

## Results

### Identification of key DNA-binding residues of BZLF1 by *in Silico* alanine scanning

To identify important residues responsible for DNA recognition by BZLF1, *in silico* alanine scanning was performed. Molecular docking between BZLF1 and DNA was carried out using pyDockDNA^28^, and the 3D structure was visualised as given in **Figure 1A**. The key residues (R179A, K181A, N182A, R183A, S186A, R187A, K188A, C189A, R190A, K192A, F193A and K194A) of BZLF1 involved in the DNA binding were identified using UCSF Chimera. To evaluate their contribution in DNA binding, each of these residues was individually substituted with alanine using Discovery Studio and the resulting changes in binding energy for BZLF1–DNA interaction were analysed **(Figure 1B)**. A significant increase in binding energy was observed for the mutants R179A, R183A, R187A, K188A, R190A, K192A and K194A compared with the wild type protein, indicating reduced DNA-binding affinity. The results suggest that these residues play an important role in stabilising the BZLF1-DNA complex and are functionally important for BZLF1 for DNA recognition and binding.

**Figure 1.**
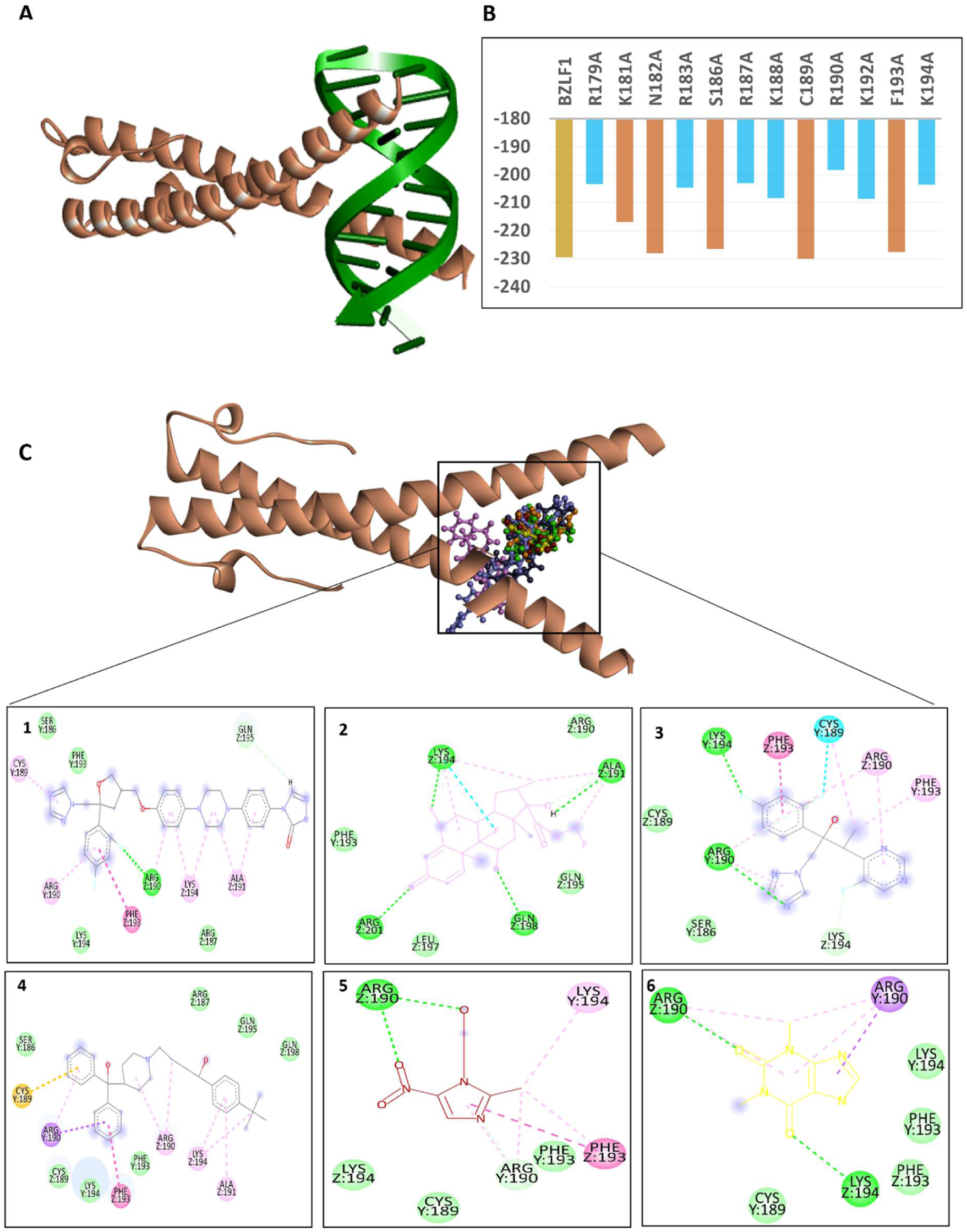
Alanine scanning and molecular docking analysis of selected drug targeting the DNA-binding domain of BZLF1. **(A)** Three-dimensional structure of the BZLF1 protein bound to DNA, highlighting the DNA-binding interface. **(B)** In silico alanine scanning key DNA-binding residues of BZLF1. Docking scores of alanine-substituted mutants of key DNA-binding residues of BZLF1 compared with the wild-type protein, showing higher impact of R179, R183, R187, K188, R190, K192 and K194 on binding affinity. **(C)** Representative 3D binding pose and corresponding 2D interaction maps (panels 1–6) of selected drugs within the DNA-binding domain of BZLF1, illustrating key molecular interactions with the identified critical residues. (1- Posconazole, 2-Fluticasone, 3- Voriconazole, 4-Terfenadine, 5 Metronidazole, 6-Theophyline).

### *In Silico* prediction of potential Anti-EBV drugs and molecular docking analysis against BZLF1

Potential anti-EBV drugs were identified using two different approaches. In-house AI/ML-based model (Anti-EBV) and miRNA-seq and RNA-seq analyses were used to identify potential anti-EBV compounds. Interactions of selected compounds with the key EBV lytic protein BZLF1 were assessed by molecular docking, focusing on the BZLF1 DNA-binding pocket to evaluate the ability of the predicted compounds to interfere with promoter recognition and viral reactivation. A total of seven compounds were identified to bind within the active site of BZLF1 **(Figure 1C)**, including Posaconazole (–7.4 kcal/mol), Terfenadine (–7.0 kcal/mol), Fluticasone (–5.6 kcal/mol), Voriconazole (–5.9 kcal/mol), Dihydroartemisinin (DHA; –5.5 kcal/mol), Theophylline (–4.3 kcal/mol), and Metronidazole (–4.2 kcal/mol). A comprehensive summary of docking scores, predicted pIC₅₀ values, and interaction details for all compounds is provided in **Table 1**.

**Table 1.**
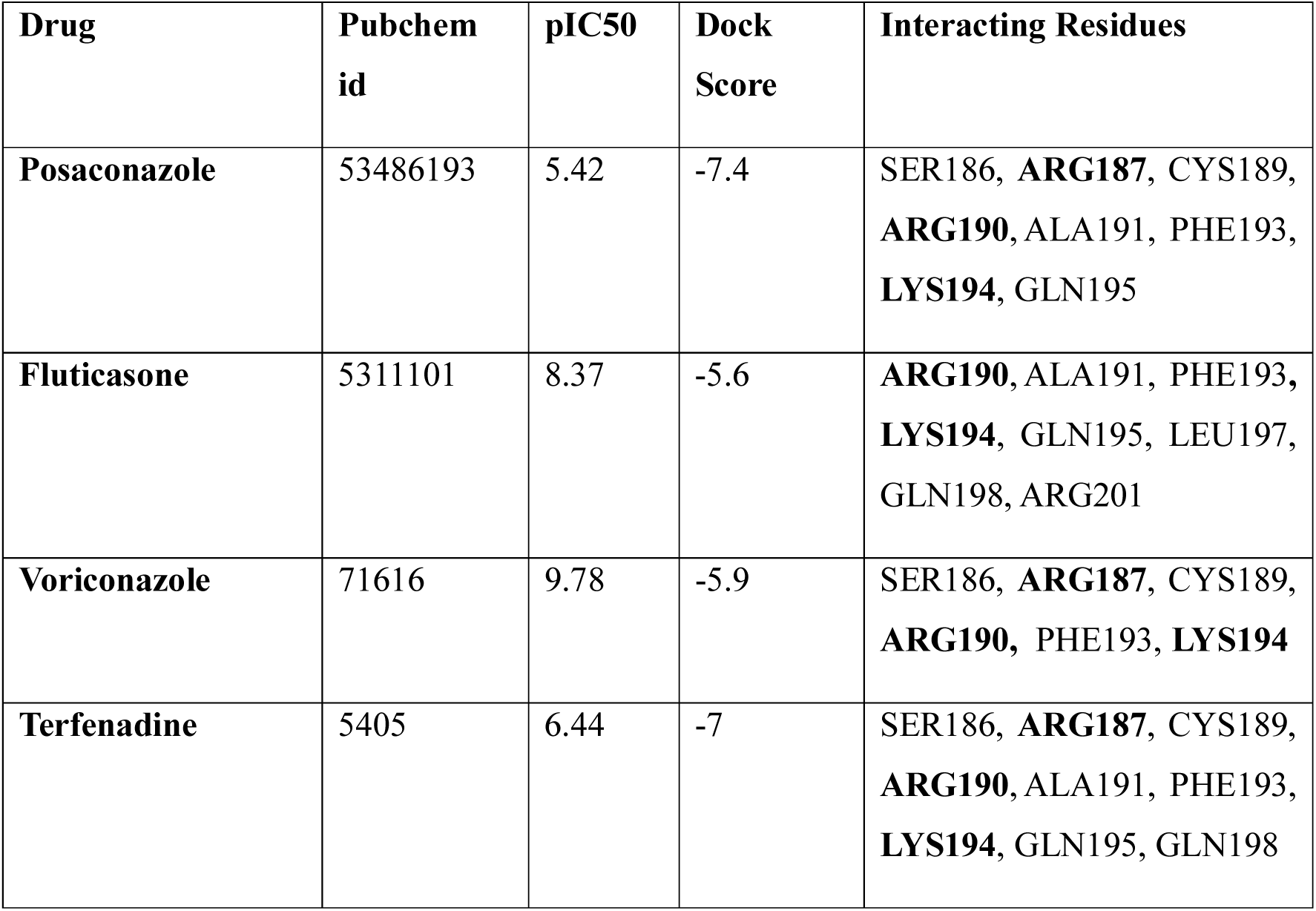
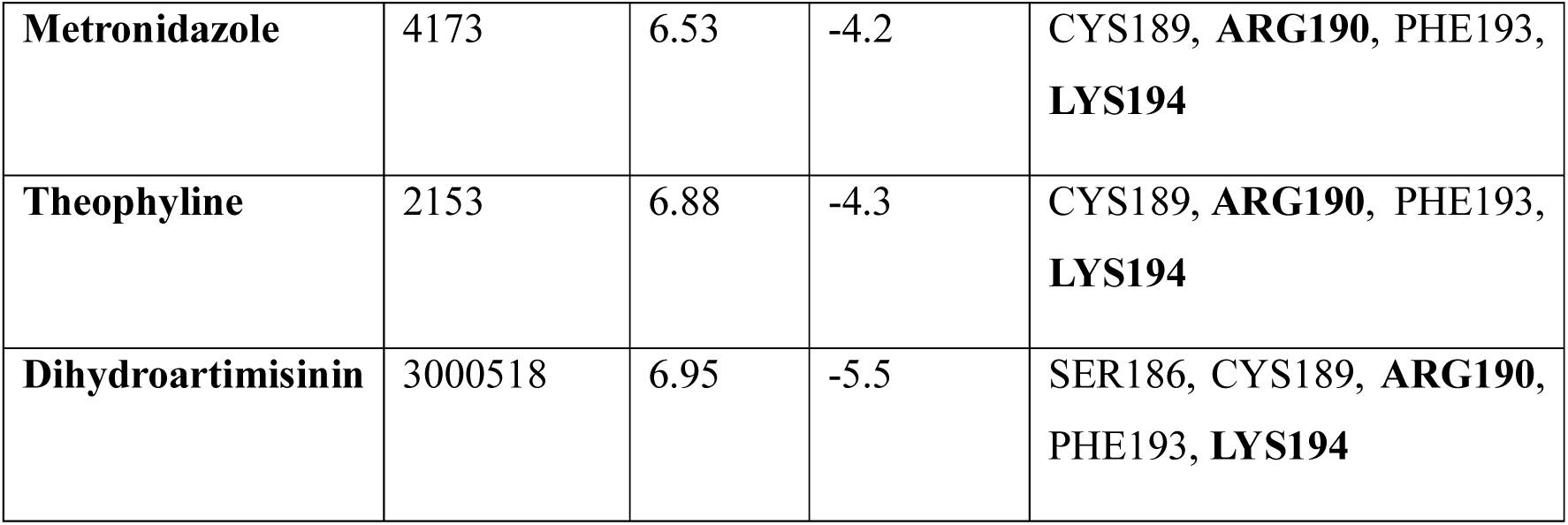
Predicted pIC₅₀ values of selected drug candidates obtained from the Anti-EBV webserver, along with their docking scores against the BZLF1 protein and corresponding interacting residues within the DNA-binding domain (Critical residues are highlighted in bold).

### In vitro screening identifies DHA as a potent inhibitor of EBV lytic genes

BZLF1 upon activation, increasess the expression of both the BZLF1 and BRLF1 genes through promoter binding. Compounds showing favourable binding affinity towards BZLF1 were further evaluated for their inhibitory potential against EBV lytic genes (BZLF1 and BRLF1) *in vitro*. Lytic reactivation of EBV was induced using sodium butyrate (SB) and 12-O-tetradecanoylphorbol-13-acetate (TPA) where SB act as deacetylase inhibitor promoting hyperacetylation at lytic genes promoter and TPA induces phosphorylation at S186 and transactivates BZLF1^29,30^. Agarose gel electrophoresis analysis confirmed that combined treatment with SB and TPA resulted in the highest induction of EBV lytic gene expression, compared to individual treatments **(Figure S1A)**. In order to identify the ideal time point for maximum EBV reactivation, the P3HR1 cells were treated with reactivating agents and examined at various time intervals (24 h, 48 h, 72 h, 96 h, and 120 h). Maximum reactivation was observed at 120 h showed by significant gene fold change of lytic genes **(Figure 2A&B)** and significant increase in viral copy number **(Figure 2C-D)**. Therefore, combined treatment (SB + TPA) at 120 h was selected for subsequent antiviral assays. For antiviral evaluation, cells were co-treated with reactivating agents and the selected seven drugs at 1 µM and 10 µM. Gancyclovir was used as a positive control which is known for its inhibitory effect on EBV replication^31^. The DNA was isolated and analysed by qRT-PCR. Gene fold change was calculated by normalizing with GAPDH using 2−ΔΔcycle threshold (Ct) method **(Figure 2E&F)**. To quantify changes in viral load, standard curve-based quantification was done. Standard curves were constructed using a plasmid vector pUC57 containing a specific insert of the BZLF1 and BRLF1 genes with a known copy number. The standard curve was generated by plotting Ct values against plasmid vector copy number **(Figure S1B&C)**. The Ct values were compared with the standard curves to determine the absolute EBV copy number **(Figure 2G&H)**. Among the tested compounds, DHA showed a significant decrease in lytic gene fold change and the copy number, indicating an inhibitory effect against lytic genes.

**Figure 2.**
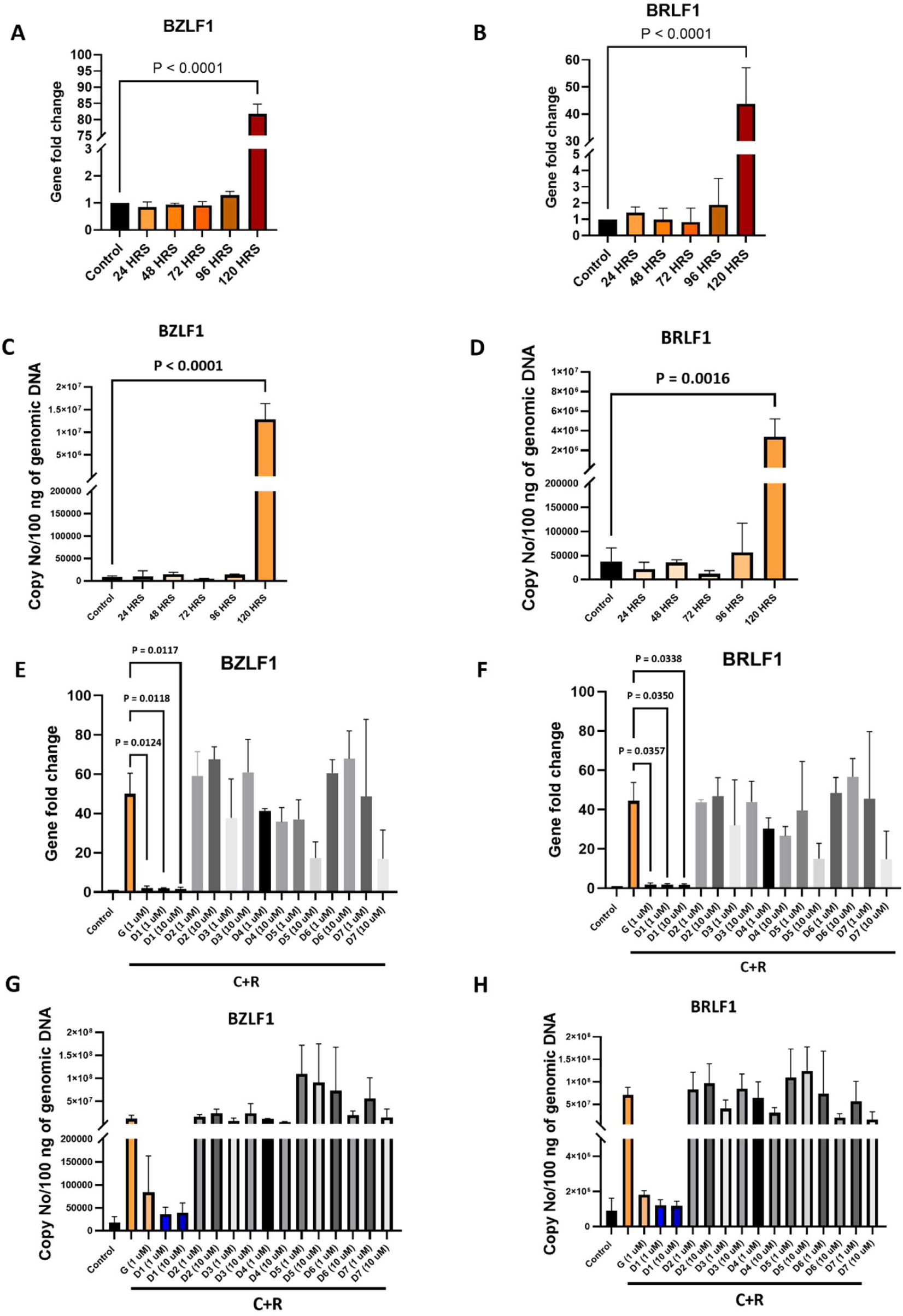
Determination of the optimal EBV reactivation conditions in P3HR1 cells and in vitro screening of candidate drugs. (A–B) Relative gene fold change of EBV immediate-early genes BZLF1 **(A)** and BRLF1 **(B)** following treatment with reactivation agents SB and TPA at 24 h, 48 h, 72 h, 96 h, and 120 h. Maximal induction of both genes can be observed at 120 h (P < 0.0001). **(C–D)** EBV DNA copy number per 100 ng of genomic DNA, quantified by targeting BZLF1 **(C)** and BRLF1 **(D)**, at the same time points after SB+TPA treatment. Viral copy number significantly increased at 120 h (BZLF1: P < 0.0001; BRLF1: P = 0.0016). **(E–H)** Effect of selected drugs on EBV lytic genes. Gene fold change of BZLF1 **(E)** and BRLF1 **(F)** in cells treated with SB+TPA in combination with selected drugs at 1 μM and 10 μM, compared to untreated control, C+R (cells + SB+TPA), and G positive control (ganciclovir, 1 μM). EBV DNA copy number per 100 ng genomic DNA following combined treatment with SB+TPA and selected drugs, measured using BZLF1 **(G)** and BRLF1 **(H)** genes. Drug treatments variably suppress EBV replication compared to C+R. Tested compounds include: D1, dihydroartemisinin (DHA); D2, fluticasone; D3, metronidazole; D4, posaconazole; D5, terfenadine; D6, theophylline; and D7, voriconazole. Data are presented as mean ± SD. DHA showed a significant decrease in lytic gene fold change and the copy number of EBV, indicating a strong inhibitory effect against lytic genes. Statistical significance is indicated as shown in the figure (P values shown). All experiments were performed in three independent biological replicates, each with technical triplicates.

### DHA inhibits BZLF1 activity through stable interactions with key DNA-binding residues

To examine the binding of DHA to residues involved in activation and DNA binding, molecular docking analysis was performed. DHA was found to interact with the S186 that inhibits transactivation as well as with two critical residues that are involved in DNA-binding inhibiting it’s binding with the oriLyt and promoter of lytic genes. This suggests that DHA may interfere with the interaction between BZLF1 and DNA, thereby inhibiting EBV reactivation and replication **(Figure 3A)**. The 3D and 2D representations of the BZLF1-DHA complex are shown in **Figure 3B**. It was observed that DHA interacted with S186, as well as R190 and K194 residues of the BZLF1 protein, which are among the key residues involved in DNA binding identified by alanine scanning. Analysis of the BZLF1 double mutant (R190A/K194A) with mutants of other critical and non-critical residues is shown in **Figure 3C**. The results revealed a weakened DNA-binding capability of the BZLF1 (R190A/K194A) mutant, highlighting the significance of these residues in the BZLF1–DNA interaction. These findings suggest that DHA may inhibit BZLF1 activity by occupying key DNA-interacting residues, thereby impairing its binding to viral promoter DNA and suppressing lytic reactivation and replication. To further investigate the effect of DHA binding on BZLF1 structural dynamics, molecular simulation studies were performed. Ligand-induced conformational changes in the BZLF1-DHA complex were analysed using CABS-FLEX 2.0^32^ **(Fig. 3D; Figure S2A)**. Most residues showed little fluctuation, indicating overall structural stability. The central region, including the DNA-binding domain, showed minimal flexibility (∼0.2–0.6 Å), while higher fluctuations (∼2–3 Å) were confined to terminal regions, consistent with their intrinsic mobility. Moreover, normal mode analysis (NMA) using iMODS^33^ showed limited deformability and stable dynamics of the complex. The low eigenvalue (3.69 × 10⁻⁵) indicated that only minimal energy was required for conformational transitions, suggesting enhanced structural stability upon ligand binding. The NMA-derived B-factor profile was consistent with experimental (PDB) pattern, and covariance analysis showed coordinated residue movements **(Figure 3E&F)**. Further, the dynamic cross-correlation matrix, deformability, and variance profile indicated that DHA binding stabilises BZLF1 without inducing major conformational changes and further supports it as a potential inhibitor of BZLF1-mediated lytic reactivation **(Figure S2B-D)**.

**Figure 3.**
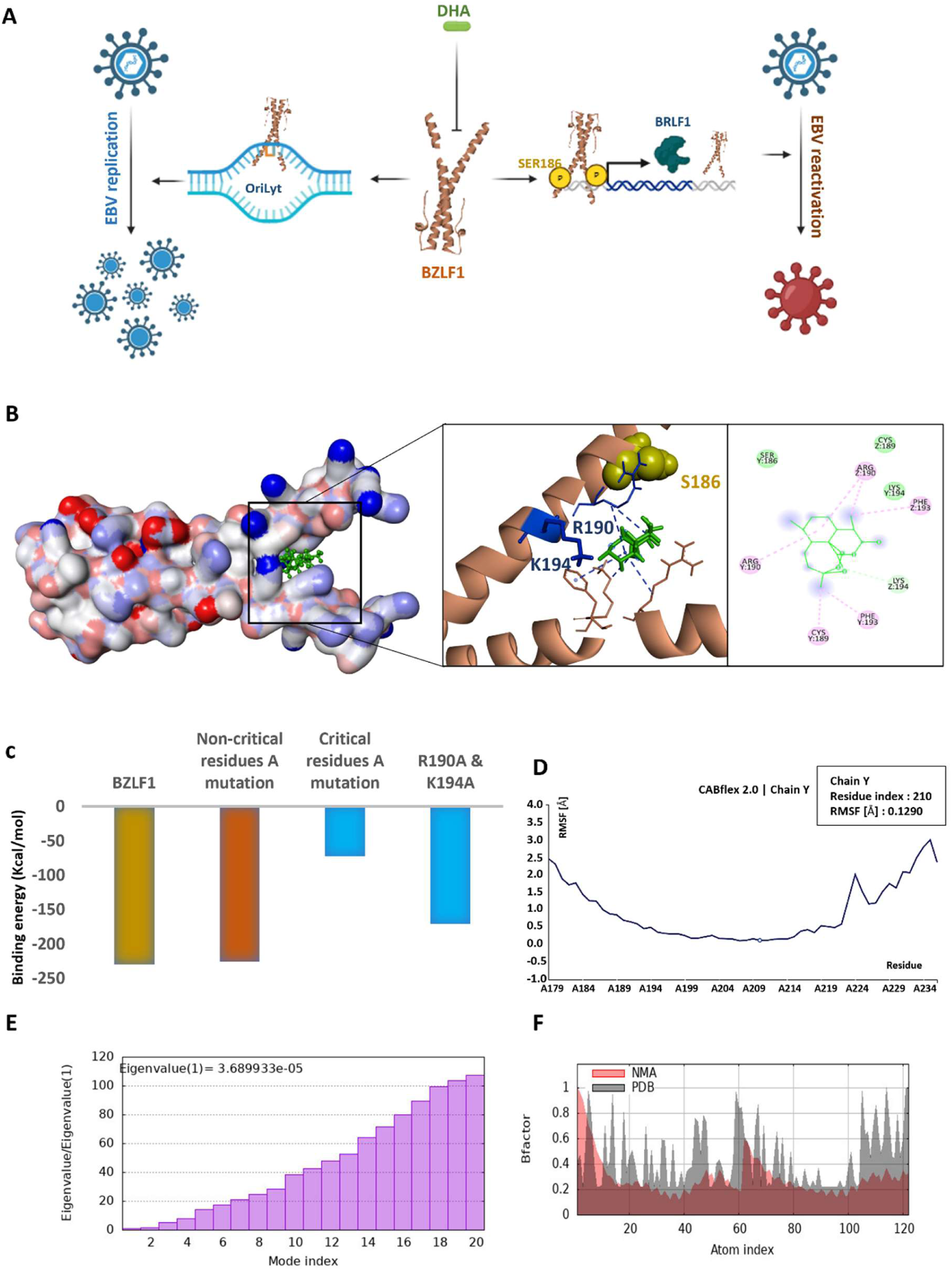
*In silico* characterization of BZLF1–ligand interaction. **(A)** Pictorial representation of DHA interacting with the S186 that inhibits transactivation as well as with two critical residues that are involved in DNA-binding, inhibiting its binding with the oriLyt and promoter of lytic genes and thus inhibiting EBV reactivation and replication. **(B)** 3-D structure of the EBV immediate-early protein BZLF1 showing the ligand-binding pocket. The structure highlights the docked ligand (green) within the binding cavity and its interaction network with surrounding amino acid residues (BZLF1). Key interactions include hydrogen bonds, carbon–hydrogen bonds, alkyl, and π-alkyl interactions with residues such as Arg190, Lys194, Cys189, Ser186, and Phe193 (Arg190 and Lys194 are the critical residues). **(C)** Binding energy comparison for wild-type BZLF1, non-critical residue mutations, critical residue mutations, and specific double mutations of DHA interacting critical residues (R190A/K194A), demonstrating the contribution of Arg190 and Lys194 residues to ligand binding stability. **(D)** Residue-wise flexibility profile of BZLF1, indicating regions of higher and lower structural fluctuation, with increased mobility observed toward terminal regions. The central region, including the DNA-binding domain, showed minimal flexibility (∼0.2–0.6 Å), while higher fluctuations (∼2–3 Å) were confined to terminal regions, consistent with their intrinsic mobility. **(E)** Eigenvalue distribution of normal modes, where lower eigenvalues correspond to easier deformability and higher flexibility of the protein structure. **(F)** Comparative B-factor analysis derived from NMA and experimental (PDB) data, indicating consistency in predicted and observed flexibility patterns across the protein.

### DHA inhibits EBV lytic replication by reducing viral gene copy number

After the initial screening, DHA was selected for further evaluation of its inhibitory activity against EBV reactivation and replication. To assess its antiviral efficacy, DHA was serially diluted (10 µM, 3.33 µM, 1.11 µM, 0.37 µM, and 0.123 µM) and P3HR1 cells were co-treated with reactivating agents (SB+TPA). After treatment, EBV reactivation was evaluated by quantifying lytic gene fold change (BZLF1 and BRLF1) normalizing with GAPDH and viral copy number using qRT-PCR analysis **(Figure 4A-D)**. A dose-dependent reduction in EBV copy number and gene fold change was observed with increasing DHA concentration. Ganciclovir was used as positive control. Further percentage inhibition was calculated relative to the reactivated control, and inhibitory concentrations were determined. DHA exhibited a 50% inhibitory concentration (IC₅₀) of 0.2015 µM for BZLF1 and 0.1534 µM for BRLF1 **(Figure S3A&B)**. Moreover, approximately 99% inhibition (IC₉₉) of EBV reactivation was achieved at 1 µM concentration, indicating strong antiviral activity. To evaluate cytotoxicity, an MTT assay was performed on P3HR1 cells indicating the CC₅₀ of DHA to be 22.7 μM **(Figure S3C)**. Based on the IC_50_ and CC_50_ values, the selectivity index (SI) of DHA was calculated to be 113.5, demonstrating a highly favourable safety and efficacy profile. In addition, the ADMET properties of DHA were analysed using pkCSM^34^. The analysis predicted favourable pharmacokinetic characteristics, including optimal lipophilicity, structural stability, water solubility, and high gastrointestinal absorption **(Figure S3D)**. These results highlight DHA as a safe and potent inhibitor of EBV lytic gene replication in a dose-dependent manner.

**Figure 4.**
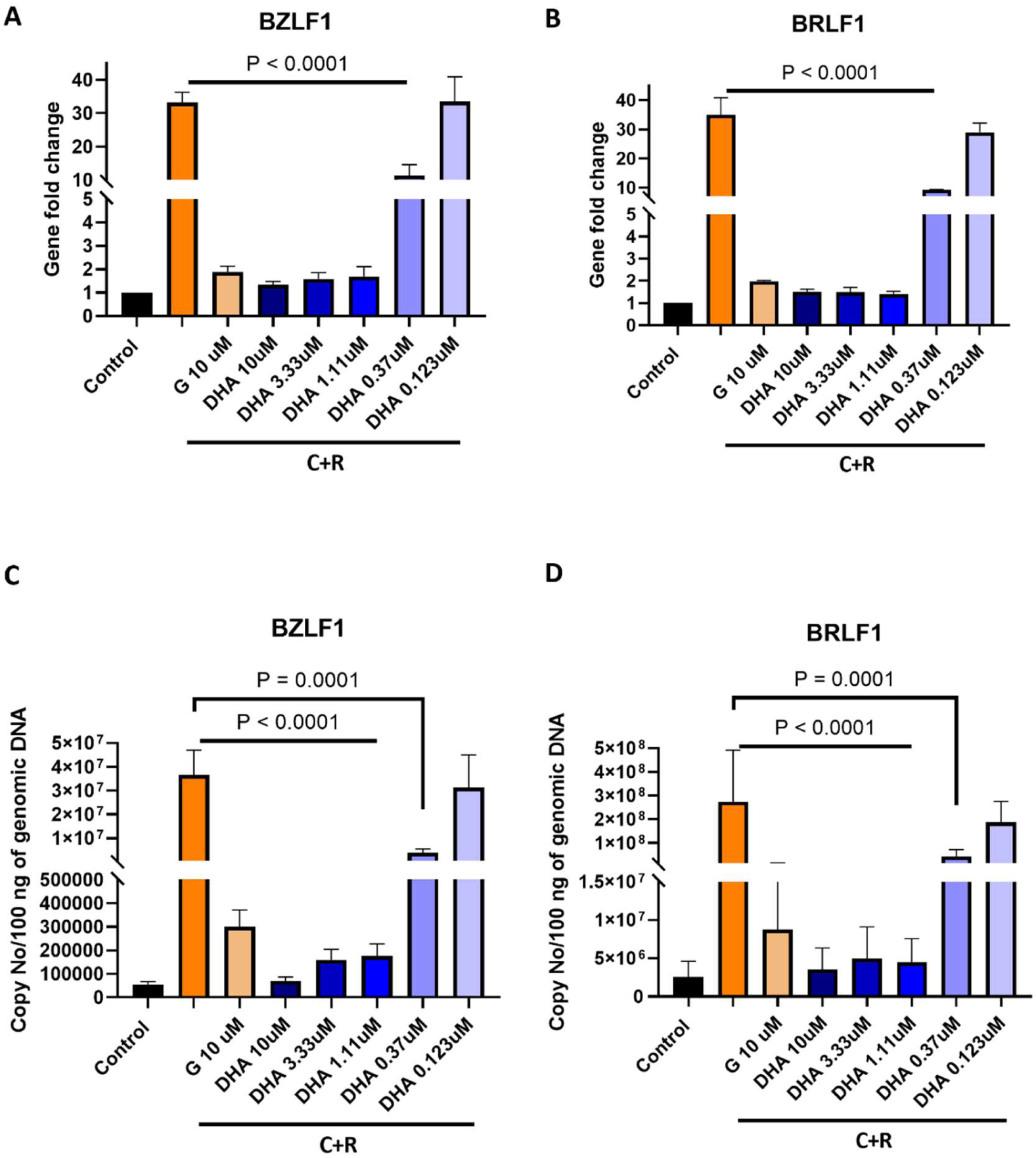
Dose-dependent effect of DHA on EBV lytic genes in P3HR1 cells. (A–B) Relative gene fold change of EBV immediate-early genes BZLF1 **(A)** and BRLF1 **(B)** following treatment with reactivation agents (C+R; SB and TPA) in combination with varying concentrations of DHA (DHA; 10 μM, 3.33 μM, 1.11 μM, 0.37 μM, and 0.123 μM). Untreated cells (control) and cells treated with the positive control drug ganciclovir (G: 10 μM) are shown for comparison. **(C–D)** EBV DNA copy number per 100 ng of genomic DNA quantified using BZLF1 **(C)** and BRLF1 **(D)** targets under the same treatment conditions. A dose-dependent reduction in EBV copy number and gene fold change was observed with increasing DHA concentration. Data are presented as mean ± SD. Statistical significance is indicated as shown in the figure(P values shown). All experiments were performed in three independent biological replicates, each with technical triplicates.

### DHA impairs early EBV lytic proteins expression associated with viral reactivation

EBV lytic gene mRNA expression was examined by qRT-PCR following treatment with reactivating agents (SB + TPA), and different concentrations of DHA. qRT-PCR analysis demonstrated a significant dose-dependent reduction in the expression of BZLF1 and BRLF1 transcripts in DHA-treated cells **(Figure 5A&B)**. These findings indicate that DHA effectively suppresses EBV lytic gene expression at the transcriptional level. To further confirm the qRT-PCR results at protein level, western blot analysis was performed. Treatment with reactivating agents increased the expression of BZLF1 and BRLF1 proteins. DHA treatment at 1 µM, corresponding to the IC₉₉ concentration, reduced expression of BZLF1 and BRLF1 proteins compared to untreated cells **(Figure 5C)**. The inhibitory activity of the positive control drug ganciclovir at 1 µM has not been observed as ganciclovir is known to inhibit viral DNA replication, but it does not effectively block EBV reactivation^31^. Dose-dependent analysis of DHA treatment was performed by using 0.2 µM, 0.6 µM, and 1 µM concentrations of DHA **(Figure 5D)**. The results showed a progressive decrease in the protein expression of BZLF1 and BRLF1 with increasing DHA concentration.

**Figure 5.**
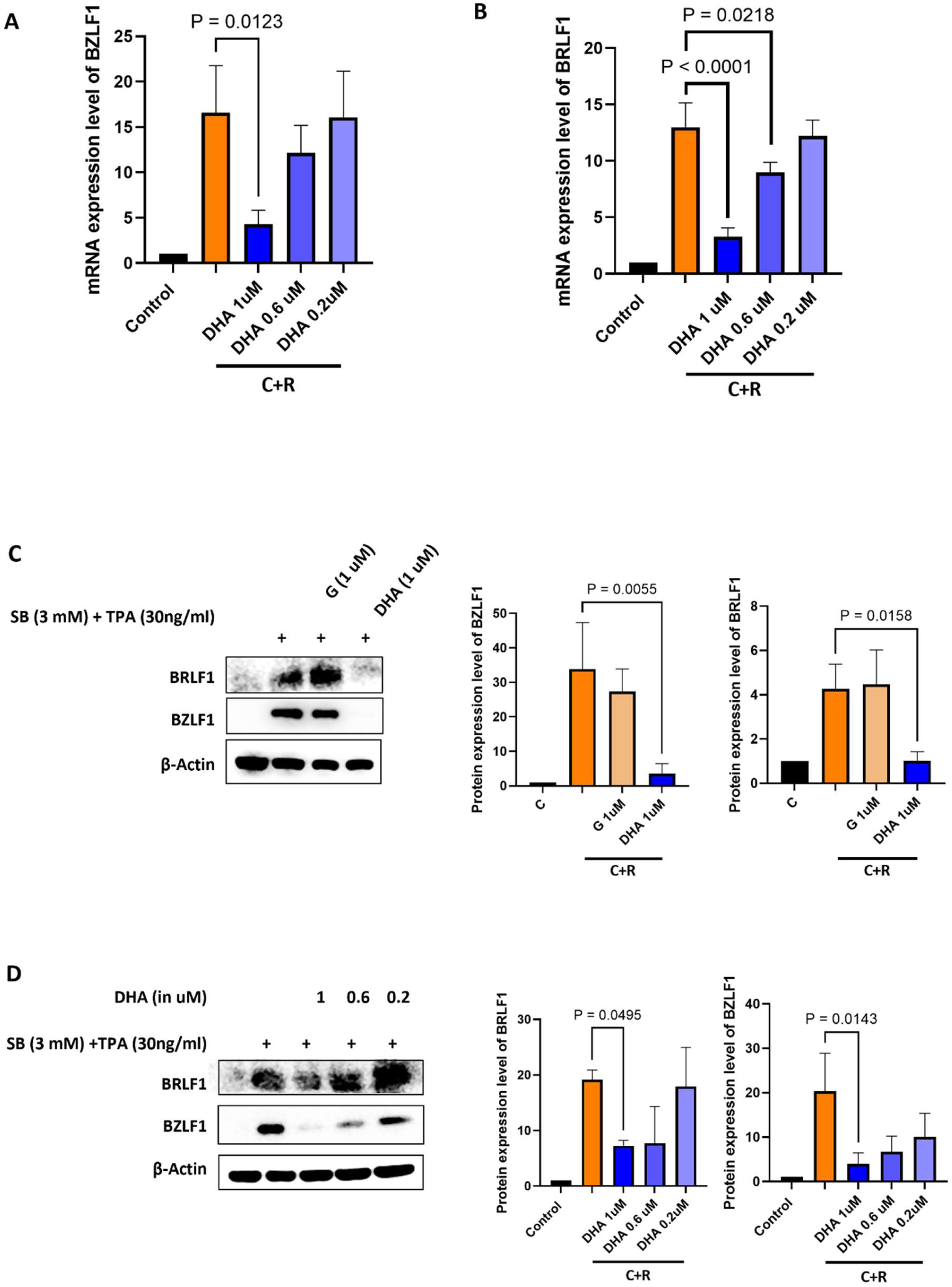
DHA inhibits immediate-early EBV lytic gene expression. (A–B) Quantitative RT-PCR analysis of BZLF1 **(A)** and BRLF1 **(B)** mRNA expression levels in cells treated with decreasing concentrations of DHA (1, 0.6, and 0.2 μM) indicating that DHA effectively suppresses EBV lytic gene expression at the transcriptional level in a dose dependent manner. **(C)** Western blot analysis of BRLF1 and BZLF1 protein expression in cells treated with SB and TPA, DHA (1 μM) corresponding to the IC₉₉ concentration and Ganciclovir (G; 1 μM), showing reduced expression of BZLF1 and BRLF1 proteins. The inhibitory activity of the positive control drug ganciclovir at 1 μM has not been observed as ganciclovir is known to inhibit viral DNA replication, but it does not effectively block EBV reactivation. **(D)** Western blot analysis of BRLF1 and BZLF1 protein expression in cells treated with SB and TPA in the presence of decreasing concentrations of DHA (1, 0.6, and 0.2 μM). Corresponding densitometric quantification of BZLF1 and BRLF1 protein levels is shown on the right highlighting a progressive decrease in the protein expression of BZLF1 and BRLF1 with increasing DHA concentration. Data are presented as mean ± SD. Statistical significance is indicated (P values shown). All experiments were performed in three independent biological replicates, each with technical triplicates.

### DHA inhibits late lytic protein expression required for virion assembly

To assess the role of DHA in inhibiting the late lytic protein, immunofluorescence was performed to analyse the expression of the EBV envelope glycoprotein gp350 in B95.8 cells. gp350 is a major EBV envelope glycoprotein involved in viral attachment and virion maturation and serves as an important marker of productive lytic replication. To determine the optimal time-point for lytic induction of EBV in B95.8 cells, latent EBV was reactivated using SB and TPA. EBV lytic induction was measured by BZLF1 gene fold change and copy number after 24, 48, 72 h by analysing lytic gene expression levels. EBV lytic induction was observed at 72 h and was therefore selected as the optimal time point for further analysis **(Figure 6A&B)**. Significantly higher gp350 fluorescence was observed in cells treated with SB and TPA, indicating successful reactivation of EBV to the lytic stage. However, a considerable decrease in gp350 fluorescence intensity was observed in cells treated with DHA compared with reactivated cells and those treated with ganciclovir **(Figure 6C)**. This demonstrates that DHA blocks the production of the late lytic protein, thereby preventing virion assembly and lytic reactivation, thereby overcoming the limitations of drugs targeting only DNA replication.

**Figure 6.**
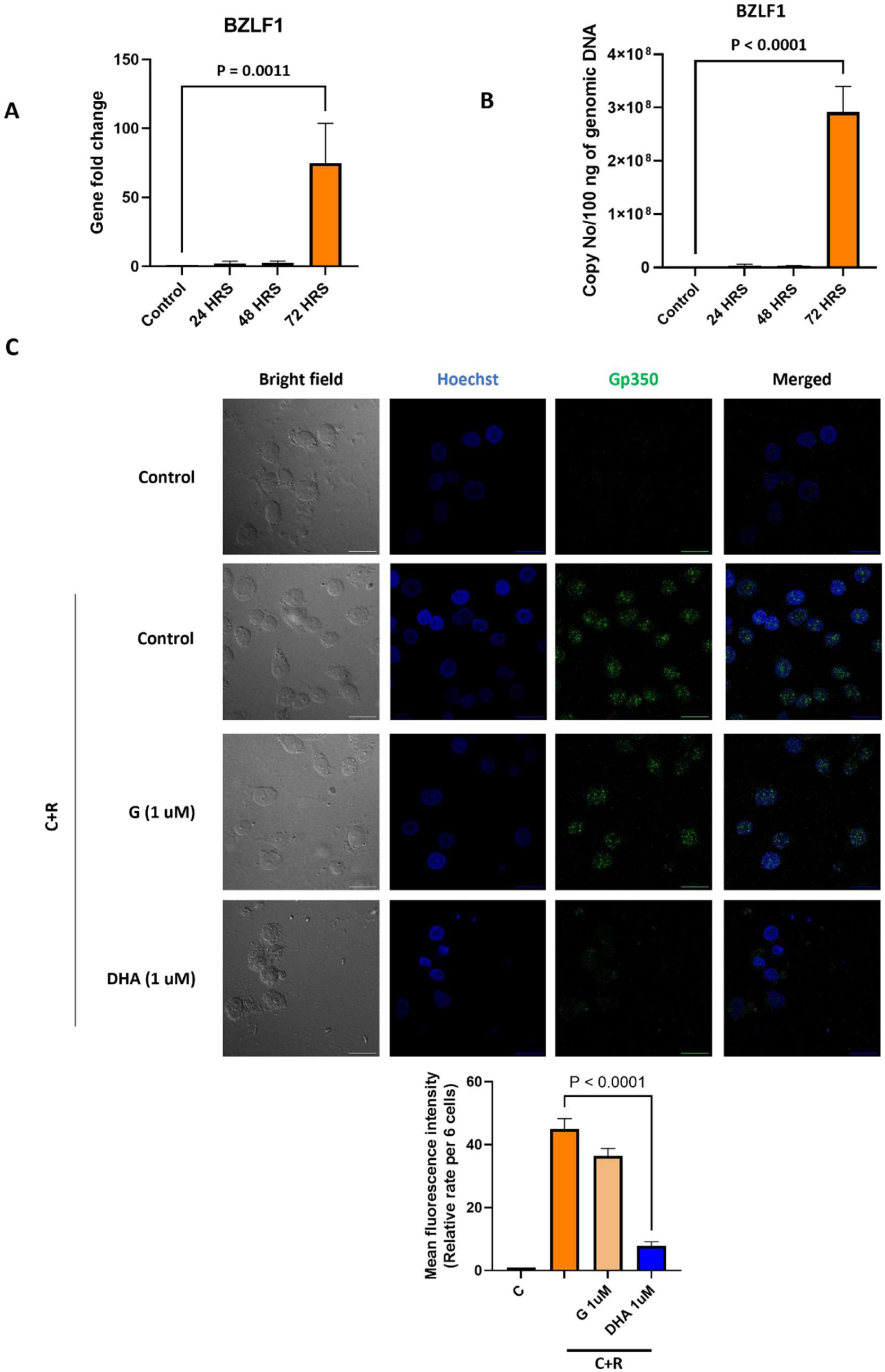
DHA inhibits late lytic protein expression in B95.8 cells. **(A)** Quantitative RT-PCR analysis showing BZLF1 gene fold change at 24 h, 48 h, and 72 h post-reactivation in B95.8. A significant increase in BZLF1 gene fold change was observed at 72 h. **(B)** Quantification of EBV genomic DNA copy number at 24 h, 48 h, and 72 h post-reactivation. A marked increase in viral BZLF1 was detected at 72 h. **(C)** Immunofluorescence analysis of late EBV lytic protein gp350. Cells under C+R conditions were treated with Ganciclovir (G; 1 μM) or DHA (1 μM). Representative images show bright-field, nuclear staining (Hoechst blue), and EBV glycoprotein gp350 expression (Alexa fluor 488 green), along with merged channel images. Increased gp350 expression is observed upon reactivation, while DHA treatment significantly reduces gp350 compared to G-treated and reactivated conditions. Quantification of mean fluorescence intensity (MFI) per 4 cells is shown below. DHA significantly decreases gp350 expression compared to control and Gan-treated groups. Data are presented as mean ± SD. Statistical significance is indicated (P values shown). All experiments were performed in three independent biological replicates, each with technical triplicates.

## Discussion

EBV is an oncogenic virus that persists in latent phase in B cells and epithelial cells. Initial infection is usually asymptomatic, but reactivation of the virus can lead to the progression of many different malignancies and non-malignant conditions. This includes nasopharyngeal carcinoma, gastric carcinomas, Burkitt lymphoma, and infectious mononucleosis. It has been observed that, in addition to latent infection, oncogenesis is also caused by lytic infection. Previous studies have demonstrated that EBV mutants defective for lytic replication have reduced capacity to cause lymphoproliferative disease, partly due to reduced levels of IL-10 and IL-6 (tumour-inducing cytokines)^35^. Moreover, studies using humanized mouse models have verified the role of EBV lytic replication in tumour development^36^. Lytic infection is also important for disease progression and the spreading of the virus. These observations highlight the importance of targeting lytic reactivation as a therapeutic strategy.

Among EBV lytic proteins, the immediate-early transactivator BZLF1 acts as a key regulator of the latent to lytic switch. BZLF1 binds to the lytic gene promoter after phosphorylation at S186, activating lytic cycle and promotes viral replication through its interaction with oriLyt. Therefore, targeting the key immediate early protein BZLF1 would be an effective strategy to inhibit EBV reactivation and replication. Several studies have explored antiviral compounds targeting different stages of the EBV life cycle, including viral DNA replication, latent gene expression, and lytic reactivation. For example, Yan et al. demonstrated that Simiao Qingwen Baidu decoction (SQBD) can inhibit cell viability and induce apoptosis in EBV-transformed B cells by modulating latency-associated genes, while also inhibiting viral replication^37^. In another study, Dipyridamole was shown to inhibit the reactivation of the virus by blocking nucleoside uptake, thereby reducing the availability of nucleotides required for viral genome synthesis^21^. In addition, Tenofovir and its derivatives, Tenofovir Alafenamide (TAF) and Tenofovir Disoproxil Fumarate, have been shown to effectively inhibit the viral DNA polymerase. TAF was also shown to be more potent and to have a longer antiviral effect^38^. However, there are limited studies that target the lytic proteins’ activity. The traditional antiviral drug discovery is often costly, time-consuming, and has a high failure rate. Therefore, drug repurposing is a promising alternative approach to traditional drug development methods, as it has known safety and pharmacokinetic data that help in accelerating the identification of effective therapeutics.

In this endeavour, we used target-based therapy, integrating both *in silico* and experimental techniques to identify the potential anti-EBV drug. We have used the in-house AI/ML-based ‘Anti-EBV’ predictive model and miRNA-Seq and RNA-Seq transcriptomics analyses to screen for potential anti-EBV compounds from the DrugBank database. BZLF1 plays an essential role in the transcriptional activation of lytic genes by binding to promoter regions, thereby triggering lytic reactivation, and also promotes viral replication through its interaction with oriLyt. Therefore, we targeted BZLF1 and screened drugs targeting its DNA-binding motif to inhibit its functional activity. Critical residues were identified by *in silico* alanine scanning, and molecular docking revealed seven compounds with favorable binding to the BZLF1 DNA-binding region. Antiviral potential of these seven compounds was further investigated by *in vitro* screening using EBV-positive P3HR1 cells. SB and TPA were used to induce lytic reactivation of EBV in the cells, where SB acts as a deacetylase inhibitor promoting hyperacetylation at the lytic genes promoter, and TPA induces phosphorylation at S186 and transactivates BZLF1^29,30^. Ganciclovir was used as the positive control for the replication study due to its known inhibitory effect on the virus’s lytic DNA replication in host cells^31^. qRT-PCR analysis of the isolated DNA after treatment with the selected compounds showed that DHA, an antimalarial drug, had the strongest inhibitory effect on EBV reactivation and replication. Docking and simulation studies further revealed that DHA forms a stable complex with BZLF1 and it interacts with S186 and two of the critical residues involved in DNA-binding that may inhibit the BZLF1 transactivation and binding with the promoter and oriLyt. The rationale was to inhibit the activation and DNA-binding ability of early lytic EBV protein BZLF1, which is critically involved in viral reactivation and replication, thereby representing a promising target-based therapeutic approach to prevent EBV infection.

Further, in vitro analysis showed that DHA was a safe and potent inhibitor of EBV lytic reactivation and replication. Analysis at the gene level confirmed its inhibitory effect on EBV replication, while expression at the transcriptional and protein levels, along with immunofluorescence analysis, indicated its inhibitory effect on EBV reactivation and virion assembly. DHA showed strong antiviral activity with approximately 99% inhibition at 1 µM. Where Ganciclovir is only effective in inhibiting DNA replication and is frequently associated with significant toxicity and carries serious safety warnings, including hematologic toxicity, impaired fertility, fetal toxicity, mutagenesis, and carcinogenic potential^39^, DHA showed highly favourable pharmacokinetic properties and an SI index of 113.5. Collectively, DHA showed greater inhibitory activity than the standard antiviral drug ganciclovir, which is effective only against DNA replication, indicating its potential as an inhibitor of EBV reactivation and replication.

In conclusion, the study demonstrates the effectiveness of integrating AI/ML-based screening with experimental validation for identifying potential antiviral agents targeting the lytic proteins that initiate the transition from the latent to the lytic phase. The results show the antiviral activity of DHA and support its potential as a therapeutic candidate targeting EBV lytic proteins, with possible implications for reducing the burden of EBV infection.

## Material and Methods

### Molecular docking

The top hits identified using the machine learning–based “Anti-EBV” prediction model and miRNA-Seq and RNA-Seq analysis were selected for docking analysis against two important lytic EBV proteins, BZLF1 (PDB ID – 2C9N) and BHRF1(PDB ID – 2WH6), which are required for viral reactivation and survival during the lytic cycle^40,41^. The three-dimensional structure of the selected compounds was downloaded from PubChem in SDF format, and converted to the PDBQT format by using Open Babel software^42^. The protein structure was prepared using AutoDock Tools, which included removing water molecules, adding polar hydrogen atoms, assigning Kollman charges, and saving the structure in PDBQT format. Molecular docking was performed using AutoDock Vina, with default parameters^43^. Multiple conformations were obtained for each protein-ligand complex, and the one with the lowest binding energy was considered for further analysis. The docked complexes were further analysed by using Discovery Studio Visualizer for interactional studies, and key interacting residues were identified, including hydrogen bonds, hydrophobic, and other non-covalent interactions, for the stability of ligands with the respective proteins^44^.

### *In silico* Alanine Scanning of the BZLF1 DNA-Binding Domain

*In silico* alanine scanning was conducted to find important amino acid residues involved in BZLF1 protein’s interaction with host DNA. Docking was performed between the BZLF1 protein and DNA using pyDockDNA. Then, the complex structure was further investigated by UCSF Chimera to detect important residues positioned on the interaction site of the BZLF1 protein with DNA within a zone of 5 Å. Important residues were replaced by alanine residues one by one using Discovery Studio to study the influence of each residue on DNA binding. The new complex structures were analysed by molecular docking to measure the impact of residue substitution on the BZLF1 protein’s DNA binding affinity. Moreover, the combined effect of substitution of all critical residues was calculated and compared with mutations of non-critical residues and the native protein BZLF1.

### Molecular dynamics simulation

The dynamic nature of the protein-ligand complexes was studied to investigate the stability and flexibility of the docked protein-ligand complexes. The residue-level dynamics of the docked complexes were studied by estimating the Root Mean Square Fluctuations (RMSF) through the online server CABS-flex 2.0. Moreover, to study the inherent dynamics of the docked complexes, a normal mode analysis (NMA) was performed using the online server iMODS. The NMA analysis also helped provide additional parameters such as B-factor, eigenvalues, and variance analysis of the complexes.

### Cell line and treatment

For *in vitro* validation, the EBV-positive cell line P3HR-1 (ATCC-HTB 62), B95.8 cells from the National Centre for Cell Science, Pune, India, were cultured at 37 °C in a humidified incubator with 5% CO_2_ in RPMI-1640 medium supplemented with 10% fetal bovine serum (FBS). For the induction of lytic replication of EBV, the cells were treated with sodium butyrate (SB, 3 mM; Sigma-Aldrich)^45^ and 12-O-tetradecanoylphorbol-13-acetate (TPA; 30 ng/mL Sigma-Aldrich)^46^. Lytic reactivation of EBV was induced using SB and TPA where SB act as deacetylase inhibitor promoting hyperacetylation at lytic genes promoter and TPA induces phosphorylation at S186 and transactivates BZLF1^29,30^. Dihydroartemisinin, posaconazole, fluticasone, voriconazole, terfenadine, metronidazole, and theophylline (Glr innovations) were used. All the drugs were prepared in dimethyl sulfoxide (DMSO) with a final DMSO concentration of less than 0.1%.

### Plasmids used and standard curve generation

BRLF1_pUC57 and BZLF1_pUC57 plasmids, synthesized by GCC Biotech India Pvt Ltd, were used for standard curve generation. Plasmids with known copy numbers were serially diluted and subjected to qRT-PCR using GoTaq Probe RT-qPCR Master Mix (Catalogue no. A6102, Promega Corporation, USA). A standard curve was created by plotting the Ct values for each dilution against the corresponding copy number and performing the regression analysis. This curve was further used for the quantification of EBV copy number in the experimental samples.

### Cell viability assay and ADMET analysis

For the MTT assay, MTT Cell Proliferation Assay Kit (Sigma-Aldrich) was used. Approximately 1 × 10⁴ P3HR1 cells and HEK293T cells were plated in each well of 96-well plates containing complete RPMI-1640 media with 10% FBS. These cells were incubated at 37°C for 24 h in humidified atmosphere with 5% CO₂. Following incubation, the cells were exposed to various concentrations of DHA. After treatment, the MTT Cell Proliferation Assay was performed following the kit’s manual procedure. Absorbance measurement of each sample was carried out at 570 nm in the microplate reader. Cell viability was measured as a percentage relative to untreated controls. The CC₅₀ value of DHA was determined from its dose-response curve. The selectivity index (SI) was determined by dividing CC₅₀ by the IC₅₀ value. The pkCSM prediction server was utilised to study the pharmacokinetic properties of DHA.

### DNA isolation and qRT-PCR analysis

To assess the potential anti- EBV activity of the selected compounds, a single-tube multiplex real-time PCR approach was performed. The immediate-early lytic genes, i.e., BZLF1 and BRLF1 genes, which play a crucial role in the lytic reactivation of EBV, were targeted. The EBV gene sequences were obtained from the NCBI database. The NCBI database accession number of the EBV genome was NC_007605.1, and the GeneIDs for the BRLF1 and the BZLF1 genes were 3783727 and 3783744, respectively. Multiple sequence alignment was performed using the EBV-positive strains, i.e., Raji (KF717093.1), P3HR-1 (LN827548.2), Akata (KC207813.1), and B95-8 (AJ507799.2). The conserved regions within BZLF1 and BRLF1 genes were identified and Integrated DNA Technologies PrimerQuest tool was employed for the designing of the primers and the probes ensuring optimal parameters for real-time PCR, including primer specificity, melting temperature, GC content, and amplicon size. The probes were then labelled with the appropriate fluorophore and quencher. Thereafter, the primers and probes were commercially synthesised by Eurofins Genomics. The multiplex real-time PCR approach allowed the simultaneous detection of the two genes. The reaction conditions were optimised. The primer and the probe concentrations, the annealing temperature, and the cycling conditions were optimised. Final concentrations of 500 nM for each primer and 250 nM for the fluorescent probe were used. The total cellular DNA was extracted using the Wizard Genomic DNA Purification Kit (Promega Corporation, USA). qRT-PCR analysis was performed to obtain Ct values for BZLF1 and BRLF1 across different experimental groups. GAPDH was used as an internal control to ensure equal input DNA. Amplification was performed using Promega GoTaq Probe RT-qPCR Master Mix (Catalogue no. A6102, Promega Corporation, USA) on the Bio-Rad CFX96 Touch Real-Time PCR Detection System. Ct values from experimental samples were compared with a plasmid-derived standard curve using regression analysis to estimate EBV copy number. Primer sequences are listed in **Table S1**. Initial denaturation at 95°C for 2 minutes was followed by amplification cycles at 95°C for 15 seconds and 55°C for 1 minute.

### RNA isolation and gene expression analysis

TRIzol reagent (Invitrogen 15596026) was used to isolate total RNA from P3HR-1 cells. Each qRT-PCR reaction contained One Step RT-PCR Mix (GCC Biotech, cat. no. GTOS1005-100R), 500 nM of each primer, and 250 nM fluorescent probe. The following cycling conditions were used for amplification: 10 minutes at 95 °C, followed by 40 cycles of 15 s at 95 °C and 45 s at 60 °C. GAPDH was utilised as the housekeeping gene for normalisation, and the gene expression of the EBV immediate-early genes BZLF1 and BRLF1 was analysed. For the calculation of relative gene expression levels, the 2−ΔΔCt method was used.

### Western blot analysis

To analyse protein expression, cells were lysed with RIPA buffer (Sigma-Aldrich) and protein was isolated in the supernatant after centrifugation at 13000 RPM. After separation by 12% SDS-PAGE, the separated proteins were transferred to a PVDF membrane (0.2 µm pore size; Novex, Life Technologies). To avoid nonspecific binding, the membranes were blocked with skim milk and incubated with primary antibodies overnight at 4 °C. The primary antibodies used were mouse monoclonal anti-EBV ZEBRA (BZLF1) antibody (1:1000; sc-53904, Santa Cruz Biotechnology) and rabbit polyclonal anti-BRLF1 antibody (1:1000; BS-4542R, Thermo Fisher Scientific). HRP-conjugated secondary antibodies, including Abcam’s anti-rabbit IgG (1:10,000; ab6721) and anti-mouse IgG (1:10,000; ab6789), were used. PierceTM ECL Western Blotting Substrate (Thermo Scientific) was used to visualise protein bands. Normalisation was done with β-actin, and band intensities were measured using ImageJ.

### Immunofluorescence assay

To determine the expression of the late lytic envelope glycoprotein gp350, an immunofluorescence assay was performed. B95.8 cells were cultured in RPMI-1640 media with 10% FBS, and 1 × 10⁵ cells were seeded on sterile coverslips placed in a 12-well plate. The treated cells were fixed with 4% paraformaldehyde and permeabilised with 0.01% Triton X-100. Following fixation, the cells were blocked in 5% BSA for 1 hour on a shaker. Thereafter, the cells were incubated with the gp350 primary antibody (1:200; sc-57724, Santa Cruz Biotechnology) overnight, followed by washing with PBS and incubation with an Alexa Fluor-labelled anti-mouse secondary antibody (1:600; ab150113, Abcam) for 2 h. Nuclear counterstaining of cells was performed using Hoechst (Thermo Scientific). Then, the coverslips were mounted on glass slides in mounting medium. To observe fluorescence, images were acquired with a Nikon confocal microscope using identical imaging parameters across all treatment groups. Signal intensity was quantified using ImageJ and compared with the control group that did not receive any treatment.

### Statistical analysis

Data analysis was performed using GraphPad Prism 9 (GraphPad Software, La Jolla, CA, USA). A one-way analysis of variance (ANOVA) test was used to determine the statistical significance across experimental groups. A p-value < 0.05 was regarded as statistically significant. Band intensities from Western blot experiments and immunofluorescence signal intensities were quantified using ImageJ.

## Author Contributions

Kumar Manoj: Supervision, Writing – review & editing, Resources, Project administration, Investigation, Funding acquisition. Vaidya Hiteshi: Writing – original draft, Methodology, Data curation and analysis, Drug screening, *in vitro* Experiments.

## Funding/acknowledgments

This work was supported by grant (<u>STS0038</u>) from the Council of Scientific and Industrial Research (CSIR)-Institute of Microbial Technology. Ms. Hiteshi Vaidya has been supported by the Department of Science and Technology, Senior Research Fellowship (INSPIRE - SRF), Award No. <u>IF210460</u>.

## Data availability statement

All the relevant data is provided in the manuscript.

